# Genetic profiling of soft tissue and bone tumors using SarcDBase

**DOI:** 10.64898/2026.05.13.724790

**Authors:** Valeria Difilippo, Karim H. Saba, Karin Wallander, Emelie Styring, Michaela Nathrath, Daniel Baumhoer, Felix Haglund de Flon, Karolin H. Nord

**Affiliations:** Department of Laboratory Medicine, Division of Clinical Genetics, Lund University, Lund, Sweden; Department of Oncology-Pathology, Karolinska Institutet, Stockholm, Sweden; Department of Clinical Genetics and Genomics, Karolinska University Hospital; Department of Orthopedics, Lund University, Skåne University Hospital, Lund, Sweden; Children’s Cancer Research Centre and Department of Pediatrics, Klinikum rechts der Isar, Technische Universität München, Munich, Germany; Department of Pediatric Oncology, Klinikum Kassel, Kassel, Germany; Bone Tumour Reference Centre at the Institute of Medical Genetics and Pathology, University Hospital and University of Basel, Basel, Switzerland; Basel Research Centre for Child Health; Department of Clinical Pathology and Cytology, Karolinska University Hospital, Stockholm, Sweden

## Abstract

To streamline molecular profiling of tumor biopsies, we developed SarcDBase, an openly accessible tool that extracts and interprets clinically relevant genetic alterations from next-generation sequencing data. By automatically linking identified variants to curated, user-defined reference lists, SarcDBase minimizes the need for specialized expertise and reduces the burden of manual data processing. The platform delivers detailed molecular profiles, diagnostic insights and an intuitive interface for comprehensive interpretation. SarcDBase’s performance was evaluated in a heterogeneous cohort of 204 deep-sequenced bone and soft tissue tumors. In most cases (81%), its interpretation closely matched the curated post-sequencing diagnosis. Discrepancies mainly occurred in samples lacking diagnostically informative mutations. In some instances, SarcDBase flagged rare or unexpected alterations, including previously unreported gene fusions. This highlights SarcDBase’s dual potential as both an interrogative research tool and facilitator of molecular diagnostics, especially for reclassification of diagnostically challenging tumor types.

## Introduction

Sarcomas are rare and heterogeneous malignant tumors of mesenchymal origin, arising in bone and soft tissues^1^. Despite their rarity, comprising roughly 1% of adult cancers and up to 20% of solid pediatric malignancies, their diagnostic complexity poses significant clinical challenges^1-6^. Subtyping traditionally relies on histopathological evaluation, anatomical site, and clinical presentation. However, benign lesions can mimic sarcomas morphologically, underlining the need for supplementary molecular-based diagnostic markers.

The current *WHO Classification of Soft Tissue and Bone Tumours* recognizes over 150 distinct entities, many with overlapping histological and molecular features^1^. While an increasing number of subtypes are defined by characteristic genetic alterations, a considerable fraction of tumors lacks recurrent or subtype-specific molecular signatures. This genetically ambiguous group is often characterized by widespread copy number aberrations and structural variants, making it difficult to distinguish biologically meaningful events from background noise^7-9^.

Advances in next-generation sequencing have paved the way for more precise molecular diagnostics, enabling the identification of clinically relevant alterations beyond morphological assessment alone^10^. In this context, computational tools that streamline interpretation and highlight key diagnostic biomarkers are essential for improving tumor classification and guiding patient management. To address this need, we developed SarcDBase — a freely available pipeline for creating mutational overview reports and interactive data visualization. By integrating high-resolution genomic and transcriptomic data from tumor biopsies with known biomarker information^1, 11^, SarcDBase eliminates the need for prior expertise or manual data curation, reducing bias and simplifying analysis.

## Materials and Methods

### Programming languages

SarcDBase is implemented in the Python programming language and executable via bespoke commands in Bash. Designed for versatility, it supports analysis across all tumor types. Users can tailor their approach by selecting prediction tools and supplying custom gene lists, enabling either comprehensive or targeted analysis. The execution module accepts modified VCF files as input and produces result files that can be used either to create per-case reports in HTML format or for dynamic visualization. The visualization interface, built with Python, Flask, and Plotly’s Dash^12, 13^, delivers interactive web-based exploration of user-selected data.

### Tumor material and clinical features

The performance of SarcDBase was tested on a clinically and morphologically diverse cohort of 204 suspected sarcomas (Supplementary Table 1). The cohort included 95 females and 109 males with ages at diagnosis ranging from 3 to 83 years (mean and median age of 58.6 and 62 years, respectively). All patients included in this study had been referred to the Karolinska University Hospital between January 2022 and May 2024 due to clinical suspicion of sarcoma. For each patient, immediately upon collection, freshly collected tissue samples, originating from either primary tumors or metastatic lesions, were snap-frozen and stored at –80°C until DNA and RNA extraction. Buffy coat cells obtained from normal peripheral blood samples or normal tissue were similarly stored at –80°C for subsequent DNA isolation.

### DNA and RNA extractions

Genomic DNA was extracted from 10 mg of tissue homogenized using a TissueLyser LT (Qiagen, Hilden, Germany) at 50 Hz for 7 minutes. Normal control DNA was isolated from 200 µL of buffy coat blood. Both extractions were performed using the DNeasy Blood & Tissue Kit, following the manufacturer’s protocols. For RNA extraction, tumor tissue was homogenized at 30 Hz for 3 minutes using the same TissueLyser LT, and RNA was purified using the RNeasy Plus Kit (Qiagen) according to the manufacturer’s instructions.

### Whole genome paired-end sequencing

Whole-genome paired-end sequencing was performed at a sequencing depth of >90x for tumor and 30x for normal DNA. DNA libraries were generated with Illumina TruSeq™ DNA PCR-Free library preparation kit (San Diego, CA, USA) as previously described^14^. In brief, ∼1100 ng of DNA was fragmented (Covaris E220), adaptors were ligated and purified, and libraries were quantified. Libraries were sequenced on the Illumina NovaSeq 6000 platform (2 × 150 bp), with demultiplexing performed using bcl2fastq2 v2.20. Sequencing reads were aligned against the GRCh37/hg19 build using BWA-MEM v0.7.17^15^. Somatic SNVs and indels were called with TNscope^16, 17^, structural variants with Manta v1.6.0 (RRID:SCR_022997), Delly v1.0.3 (RRID:SCR_004603), and TIDDIT v3.3.2 (RRID:SCR_024361). Copy number alterations were assessed using ascatNgs v4.5.0^18^, CNVpytor v1.2.1 (RRID:SCR_021627), and CNVkit v0.9.9^19^. The variants were further analyzed using Molecular Tumor Board Portal^20^ and Integrative Genomics Viewer^21, 22^. The genomic complexity score (GCS), previously published by us, was used to analyze the genomic complexity of individual DNA samples^23^.

### RNA sequencing data analysis

RNA sequencing was performed using the Illumina Stranded mRNA Prep kit as previously described^14^. Fusion gene calling was performed using the nf-core/rnafusion pipeline (v2.3.4), with Arriba, STAR-Fusion, and FusionCatcher employed as fusion callers, as previously described^14^.

### External sources

Outputs from single nucleotide and structural variant data were cross-referenced against genes listed in the *WHO Classification of Tumours, Soft Tissue and Bone Tumours*^*1*^. Gene fusion findings were cross-referenced with the Mitelman Database of Chromosome Aberrations and Gene Fusions in Cancer^11^.

## Ethics statement

The study was approved by the Swedish Ethical Review Authority (Dnr 2023-01550-01, Dnr 2022-05409-01, and Dnr 2013/1979-31 with amendment 2018/2124-32).

## Code availability

All components of the SarcDBase pipeline, including source code, documentation, and usage examples, are openly accessible via GitHub at https://github.com/ValeriaDifilippo/SarcDBase.

## Data availability

The raw sequencing data is not publicly available as it contains information that may compromise research participants’ anonymity.

## Results

### SarcDBase workflow and output

SarcDBase comprises two main modules: Execution and Visualization, as depicted in Figure 1A. The Execution module processes all types of detected mutations per case and cross-references them with curated gene lists from external sources, as specified above. This integration enables identification of both known pathogenic alterations and novel variants with potential biological relevance, such as previously unreported fusion gene partners. Users retain full control over database selection and gene list customization, provided input data is formatted according to SarcDBase specifications.

**Figure 1.**
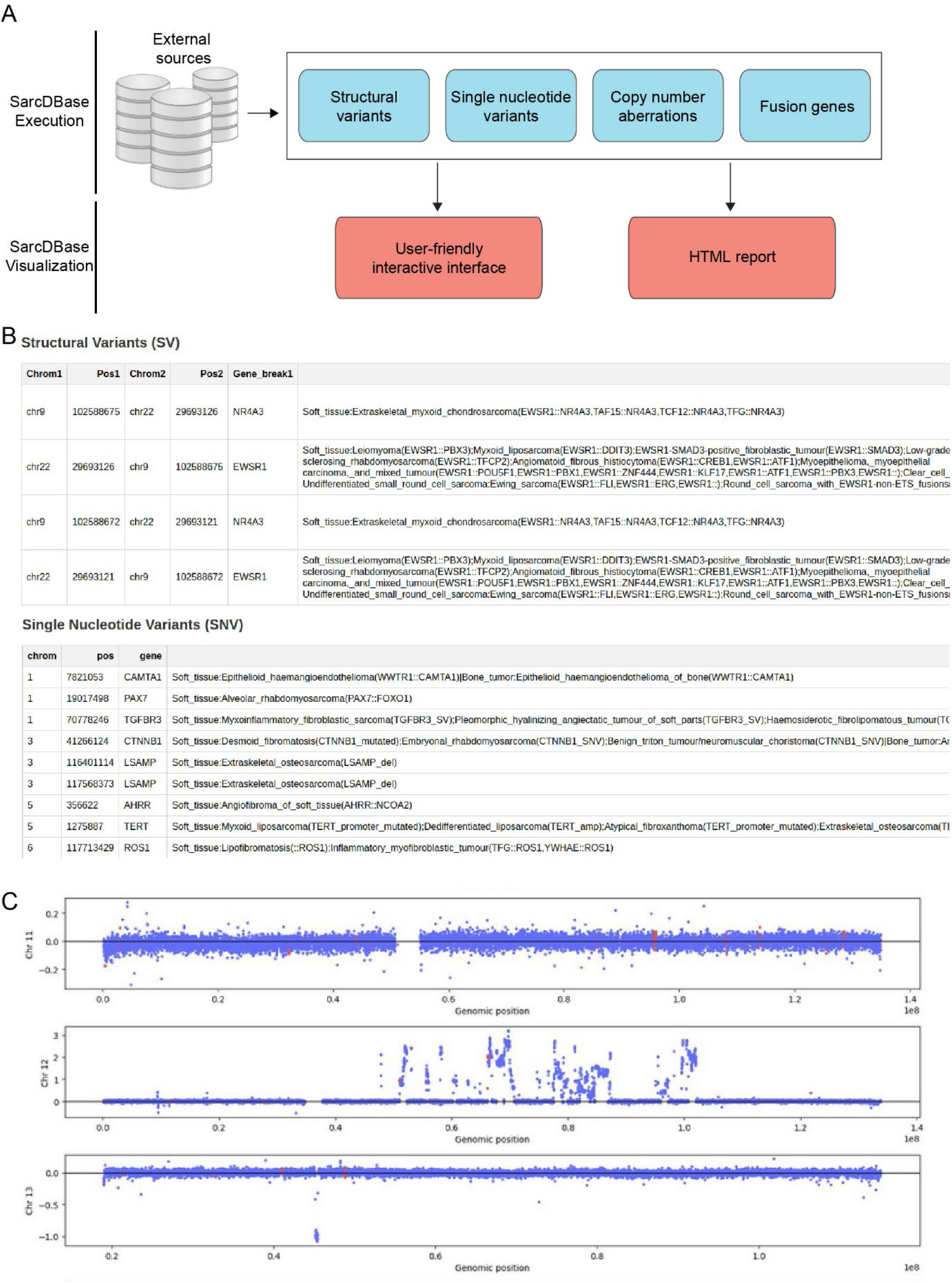
Overview of SarcDBase’s workflow and output. **(A)** SarcDBase is implemented in Bash and Python, designed to handle files in a modified VCF format. This adaptability allows it to support data from any tumor type. Users can also tailor external source inputs and integrate novel mutations, enabling cohort re-screening with enhanced flexibility and precision. **(B)** In the HTML report generated by SarcDBase, an overview of structural variants and single nucleotide variants detected in genes present in the user-provided list are shown in a tabulated format, along with possible diagnostic predictions. **(C)** Whole-genome copy number profiles are also presented per chromosome, allowing for quick scanning of large-scale copy number alterations such as chromosome arm 12q amplifications.

SarcDBase generates an HTML report summarizing all detected structural variants, single nucleotide variants, indels and genes fusions that match entries in the user-specified gene list. Genome-wide copy number alterations are visualized per chromosome, with genes from the list highlighted in red to provide a clear overview of large-genomic changes (Figure 1B-C).

The Visualization module provides a dynamic web-based interface, built to support intuitive mutation interpretation (Supplementary Videos 1 and 2). Users can explore reported variants, assess predicted diagnoses, and screen for novel mutations. Each case is displayed with both graphical and tabular summaries, and the platform allows users to survey all cases in one location, streamlining the identification of recurrent genetic events.

### Performance assessment of SarcDBase

SarcDBase was evaluated across a cohort of 204 suspected sarcomas. As previously described^14^, the genetic tests were completed after routine histology, and the impact of whole genome and transcriptome sequencing on the tumor classification was assessed for each case (referred to as pre- and post-sequencing diagnosis). In 18 cases (9%), the genetic information led to diagnostic revision (Supplementary Table 1).

The post-sequencing diagnoses were subsequently compared to those generated by SarcDBase, however, SarcDBase itself was not part of the diagnostic process. In most cases, SarcDBase aligned closely with the curated post-sequencing diagnosis, confirming known subtype-specific alterations and identifying diagnostically relevant mutations (Figure 2, Supplementary Table 1).

**Figure 2.**
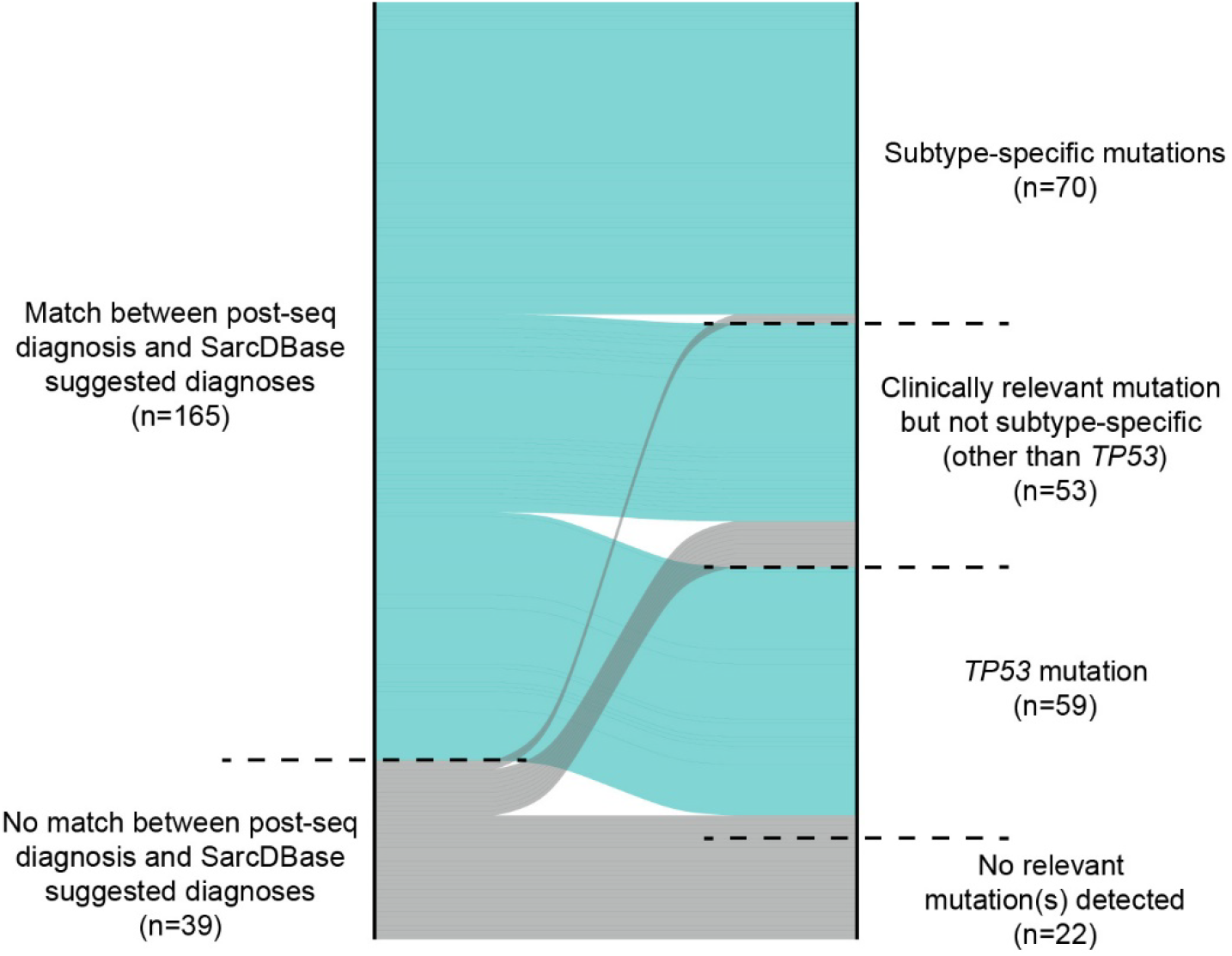
Diagnostic accuracy of SarcDBase in clinical cases. SarcDBase demonstrates strong predictive capability by providing accurate diagnostic suggestions in the majority of cases that harbor clinically relevant mutations. In these instances, the platform effectively integrates mutational profiles with curated reference datasets to propose diagnoses that align with expert clinical interpretation. Its performance highlights the value of computational augmentation in sarcoma diagnostics, particularly in complex or ambiguous cases where genomic insights can guide classification.

Among 70 cases harboring subtype-specific mutations, defined as characteristic of a tumor subtype per WHO classification^1^, SarcDBase accurately predicted the diagnosis in 68 cases (97%; Figure 2). However, there were discrepancies for two cases: in a case harboring rearrangement of the *PRKCB* gene, SarcDBase suggested deep fibrous histiocytoma, while the clinical diagnosis was lipoma; in a case positive for *EWSR1::SMAD3* fusion, SarcDBase proposed *EWSR1::SMAD3*-positive fibroblastic tumor, while the post-sequencing diagnosis was low-grade chondrosarcoma. It is important to note that SarcDBase is agnostic to clinicomorphological parameters which would normally weigh into reaching diagnoses.

In cases harboring clinically relevant but non–subtype-specific mutations (excluding *TP53*; n = 53), SarcDBase generated multiple potential diagnoses. These results require integration with clinicopathological data before a definitive diagnosis can be established. Representative mutations in this category include amplifications of *CDK4* and *MDM2*, as well as structural and single nucleotide variants in genes such as *HMGA2, RB1, CDKN2A*, and *ATM*. Although *TP53* mutations are not tumor-type specific, they are strongly associated with high-grade sarcomas characterized by extensive genomic complexity (Figure 3). In such cases (n = 59), SarcDBase appropriately generated multiple differential diagnoses. Definitive classification requires integration of both genomic findings and clinicopathological context. In the 22 remaining cases (11%), no diagnostically informative mutations were identified, limiting SarcDBase’s contribution beyond the initial clinicopathological assessment.

**Figure 3.**
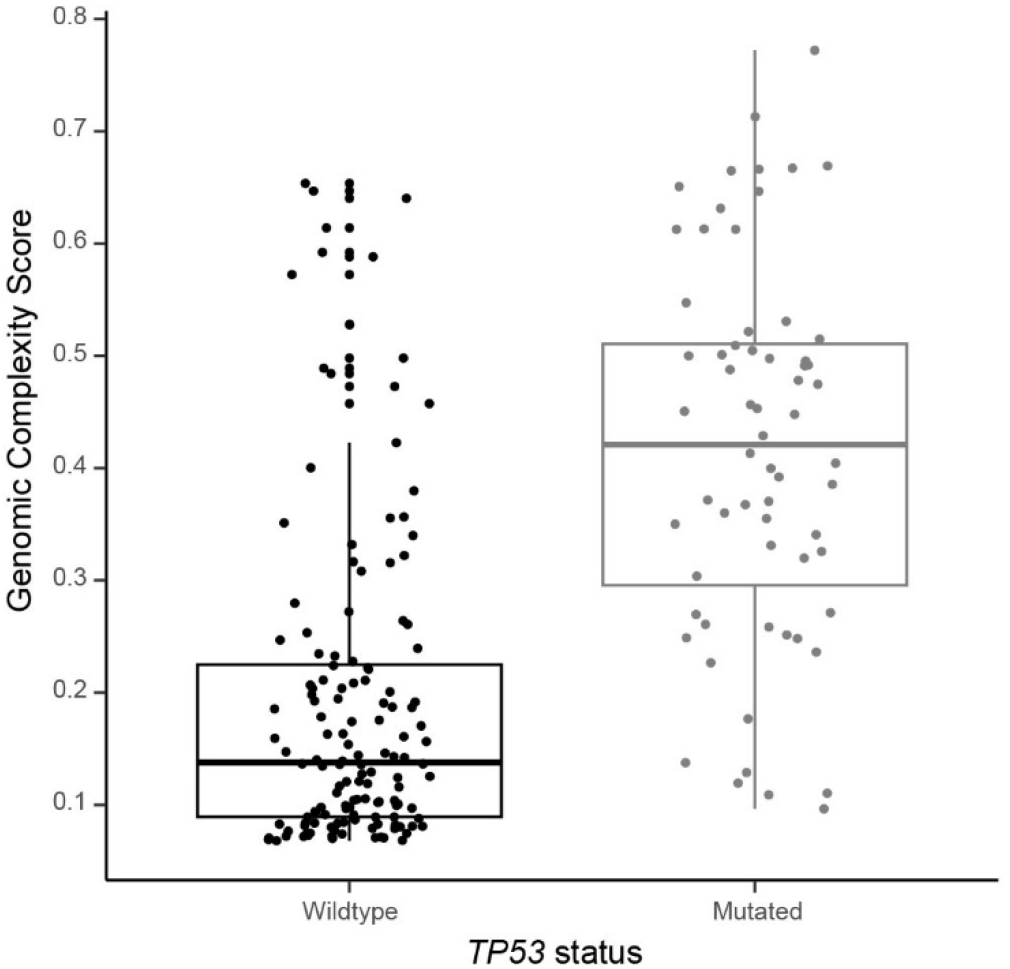
Cases with *TP53* mutations exhibit a greater degree of genomic alterations compared to *TP53* wildtype cases. The Genomic Complexity Score (GCS) was computed using the mean and standard deviation (SD) of log_2_-transformed genomic copy number values. For each autosome, a normalized GCS was calculated by averaging the absolute values at ±1.96 SD from the mean. The final GCS for each case was obtained by averaging the normalized scores across all 22 autosomes. Cases with *TP53* mutations exhibit a higher GCS compared to *TP53* wildtype cases.

## Discussion

Molecular analysis has become an essential component for sarcoma diagnostics, with subtype-specific gene fusions or other mutations serving as characteristic markers for an increasing number of subtypes^1, 24-31^. However, the growing number of such markers, combined with the widespread availability of high-resolution sequencing data in both clinical and research settings, has introduced new challenges. Chief among them is the difficulty of extracting biologically and diagnostically significant alterations, particularly in tumors with highly complex genomes. In these cases, the distinction between driver and passenger mutations becomes increasingly blurred, necessitating careful interpretation.

SarcDBase can address a critical unmet need in molecular pathology by streamlining the interpretation of complex genomic data. It highlights diagnostically relevant genetic alterations and maps them to associated disease entities, thereby facilitating the clinical interpretation of large-scale sequencing datasets. Moreover, SarcDBase is technically capable of identifying any treatment-predictive biomarker of interest, as long as it is included in the user-defined list. This empowers users to translate molecular findings into actionable therapeutic strategies and personalized treatment plans.

The platform’s main constraint lies in the gene list provided by the user. As proof of principle, we chose to restrict this list to genes included in the most recent update of the WHO classification. However, numerous new diagnostic, prognostic, and potentially predictive biomarkers have been identified since that update, underscoring the importance of regularly expanding and refining the gene list to keep pace with emerging discoveries. SarcDBase was designed to address this challenge by providing a comprehensive analysis of the mutational profile across individual tumor cases. It screens for single nucleotide variants, indels and structural alterations, plots copy number changes and automatically matches with established biomarkers from curated external sources. This unsupervised approach minimizes the need for manual data filtering and prior expert knowledge, thereby reducing bias and enabling efficient interpretation. SarcDBase is compatible with matched DNA-RNA datasets, as well as DNA or RNA alone, offering flexibility across different sequencing workflows.

In the current evaluation, SarcDBase successfully identified subtype specific diagnostic biomarkers and suggested the correct diagnosis in 165 out of 204 cases (81%), while 39 cases (19%) lacked sufficient molecular evidence to support a distinct diagnosis. The tool reliably identified well-established diagnostic markers including, but not limited to, the *SS18::SSX1/2, NAB2::STAT6, EWSR1::NR4A3* gene fusions, and *KIT/PDGFRA* single nucleotide variants, when present^29, 32-35^. In contrast, ambiguous variants, along with copy number and structural aberrations involving *MDM2, HMGA2, RB1, CDKN2A*, and *TP53*, required integration with clinical and histopathological information to guide diagnosis.

To further enhance SarcDBase’s diagnostic and research capabilities, users can customize reference sources by incorporating newly validated markers. However, it is important to emphasize that SarcDBase is not designed to function as a standalone diagnostic tool and was not tested as such. Accurate classification of sarcomas still requires expert curation of molecular findings, integration with histopathological evaluation, and consideration of clinical and imaging data within the appropriate diagnostic or research context. One of the primary limitations of SarcDBase lies in the input data. Beyond the gene list specified by the user, the tool’s performance is closely tied to the reliability of upstream variant detection methods. Errors such as misclassification or missed events at this stage can significantly affect downstream interpretation. Despite these constraints, SarcDBase offers several key advantages. It provides an automated workflow and enables efficient visualization of large-scale genomic datasets. The platform also supports reproducible analysis pipelines, fostering consistency across research efforts. Its compatibility with diverse data formats and user-friendly interface ensures accessibility for both computational biologists and clinical researchers.

## Acknowledgements

VD was supported by the Royal Physiographic Society, Lund, Sweden. ES was supported by the Skåne University Hospital donationsfonder and Greta och Johan Kocks stiftelser. DB was supported by the Swiss National Science Foundation, the Hemmi Stiftung, the Foundation of the Basel Bone Tumour Reference Centre, the Gertrude von Meissner Stiftung, the Susy-Rückert Gedächtnis-Stiftung, the Nora van Meeuwen-Häflinger Stiftung, and the Stiftung für krebskranke Kinder, Regio Basiliensis. KW and FHdF were supported by the Swedish Cancer Society (grant IDs 19 0029 JCIA; 22 0607 FE, and 22 2124 Pj), the Swedish Sarcoma Association (Sarkomföreningen), the Cancer Society in Stockholm (Radiumhemmets forskningsfonder) and Karolinska Institutet. KHN was supported by the Swedish Childhood Cancer Fund, the Swedish Cancer Society, the Swedish Research Council, and the Swedish state under the agreement between the Swedish government and the country councils (the ALF agreement).

## Author contributions

VD, KHS, ES, and KHN conceptualized and designed the study. VD led the development of SarcDBase. Bioinformatic analyses and result interpretation were performed by VD and KHS. DB and MN provided foundational data instrumental in developing the SarcDBase prototype. KW and FHdF contributed sequencing data and relevant clinical information. FHdF conducted the histopathological review. All authors provided intellectual input. VD, KHS, and KHN drafted the manuscript with contributions from all other authors.

## Supplementary material

**Supplementary Table 1**. Cohort summary.

**Supplementary Video 1. SarcDBase structural and single nucleotide variants visualization** Users have the flexibility to navigate through multiple sections in the SarcDBase app based on data availability. For instance, genomic structural and single nucleotide variants and associated annotations can be visualized in an interactive tabular format as shown.

**Supplementary Video 2. SarcDBase copy number visualization**. The copy number section provides a visual representation of the genomic copy number profile for each individual chromosome. Users have the capability to zoom in on specific regions or genes of interest. Notably, red dots are used to highlight genes included in the user-provided list, which in this case were those mentioned in the *WHO Classification of Tumours, Soft Tissue and Bone Tumours*^*1*^.

